# ION-Decoding: A Single-channel Interactive Offline Neural Decoding Algorithm for a large number of neurons

**DOI:** 10.1101/2020.11.16.385583

**Authors:** Mohsen Rastegari, Hamid Reza Marateb

## Abstract

Researchers have widely used extracellular recordings as a technique of paramount importance due to its wide usage in cognitive studies, health technologies, and prosthetics and orthotics research. To extract the required information from this technique, a critical and crucial step, called spike sorting, must be performed on the recorded signal. By this method, it is possible to analyze a single neuron (single-unit activity) and investigate its specifications, such as the firing rates and the number of action potentials (spikes) of an individual neuron. Here we introduce a novel idea of a user-friendly interactive, offline, and unsupervised algorithm called ION-Decoding. This platform extracts and aligns the spikes using a high-resolution alignment method, and the clusters can be atomically identified and manually edited. The entire procedure is performed using the minimum number of adjustable parameters, and cluster merging was performed in a smart, intuitive way. The ION-Decoding algorithm was evaluated by a benchmark dataset, including 95 simulations of two to twenty neurons from 10 minutes simulated extracellular recordings. There was not any significant relationship between the number of missed clusters with the quality of the signal (i.e., the signal-to-noise ratio (SNR)) by controlling the number of neurons in each signal (p_value=0.103). Moreover, the number of extra clusters was not significantly dependent on the parameter SNR (p_value=0.400). The accuracy of the classification method was significantly associated with the decomposability index (DI) (p_value<0.001). A number of 77% of the neurons with the DI higher than 20 had the classification accuracy higher than 80%. The ION-Decoding algorithm significantly outperformed Wave_Clus in terms of the number of hits (p_value=0.017). However, The Wave_Clus algorithm significantly outperformed the ION-Decoding algorithm when the false-positive error (FP) was considered (p_value=0.001). The ION-Decoding is thus a promising single-channel spike sorting algorithm. However, our future focuses on the improvement of the cluster representative identification and FP error reduction.

## Introduction

The extracellular recording is one of the common ways that neuroscientists use to study and discover the brain and the electrical activities of living neurons both in vivo and in vitro. The experimenter records the extracellular space that has produced action potential around the active neuron by placing the electrode in that area. In other words, the action potential or the spike is the voltage difference between the tip of the recording electrode in the extracellular space of the neuron of interest and a ground electrode placed far from that area. The electrode tip will record the action potentials related to different neurons around the electrode by placing the electrode into the brain tissue and starting the extracellular recording. As a result, the recorded signal will be a mixture of spikes related to different neurons, and to find out and extract the activity of a single neuron, it is necessary to sort and assign these different spikes to their specific cells. This process is called spike sorting (1, 2). In other words, the spikes produced by different neurons have various shapes. These differences come from factors like the morphology of the neuron’s dendritic tree, spacing, and direction related to the recording location and physiological attributes for each neuron. Therefore, as another definition, grouping obtained spikes according to their similarity in shape into clusters is known as spike sorting. The detected clusters are related to the activity of different neurons (3).

Extracellular recording contains high-resolution information from neural tissue and reveals information from the input (synaptic activity) and output (spiking) of the neuron of interest in the recording area. Therefore, it is a powerful tool for scientists to study and explore the Central Nervous System(CNS) pathway and neural network (4). Studies such as inspecting different parts of the brain, detecting and distinguishing engaged portions of the brain during a specific task, and investigating mirror neurons, extracellular recording, and the information decoded from the single-unit activity play a crucial role (5). Moreover, scientists are recently working on discovering how extracellular recording results could help patients dealing with refractory epilepsy. They have also started to see if this type of recording could help medical technologies treat neurological diseases like paralysis, epilepsy, and cognitive and loss of memory (2, 6). Besides clinical purposes, using the extracellular recording in applications like Brain-Machine Interface (BMI) needs hardware spike sorting for detecting single-unit activity and data reduction toward wireless data transition as the technology usually depends on the single-unit activity as input. In recent studies in BMI, scientists are working on adding touch and force sensors to the prosthesis by using the extracellular recording as part of the process to get constant access to neurons’ activity (7, 8).

According to the extracellular recording’s unique properties and the extensive usage of this technique in both clinical and experimental studies, designing a smart algorithm that could precisely analyze the physiological data is essential. Because of that reason, many spikes sorting algorithms with different approaches have been designed. In a point of view, it is possible to divide all the spike sorting algorithms into three groups: some of which like Offline Sorter (Plexon, Inc.) (9) are commercial and consequently not accessible for everyone, others like OSort (10), Wave_Clus(11), KiloSort(12), etc. are open-access software packages that are common in use, and there is not any reported commercial/open-access software or package for the rest.

Apart from the implementation, the structure of the spike sorting algorithm for most of the software or packages consists of four processing steps: 1. Filtering, 2. Spike detection, 3. Alignment and feature extraction, 4. Clustering (3). This procedure was briefly shown in **Fig. 1**.

**Fig. 1.**
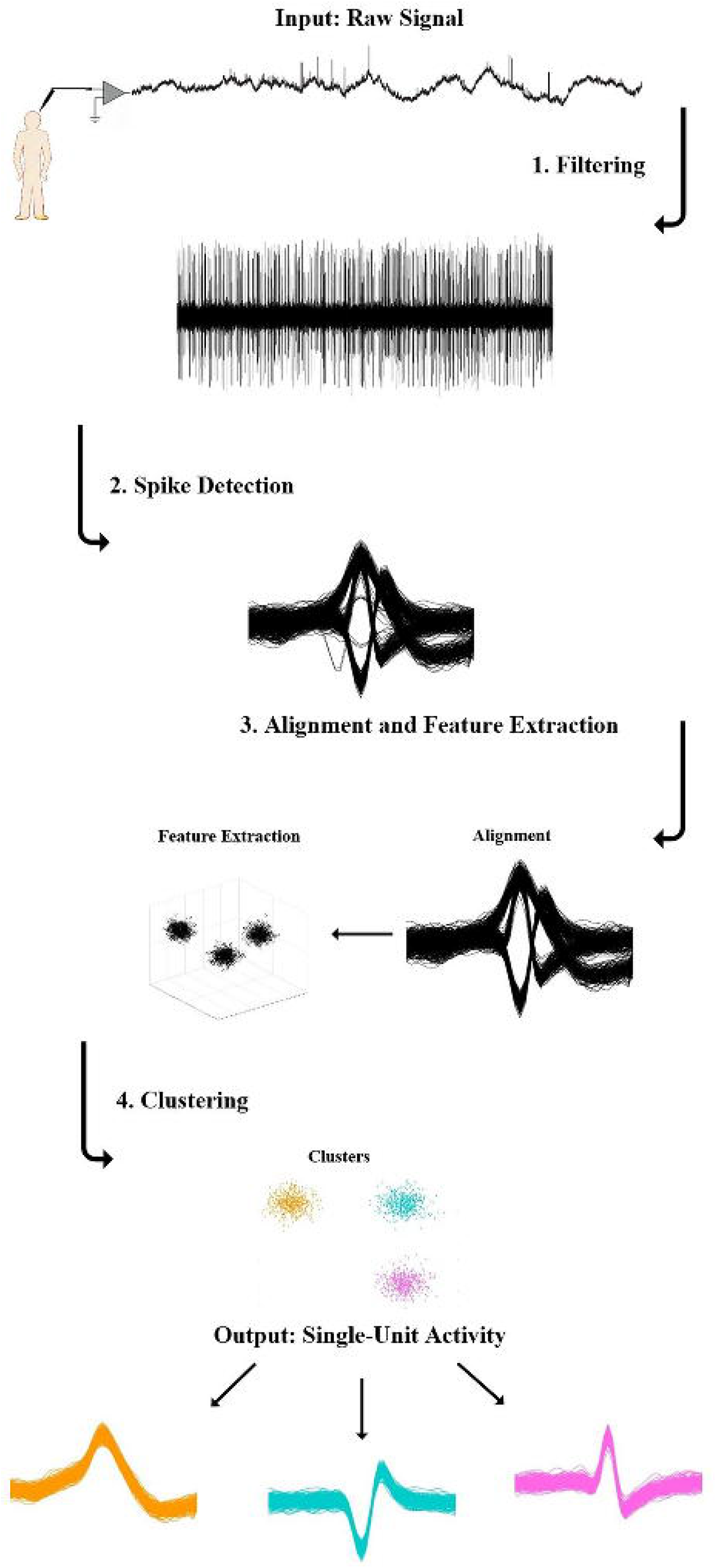
Common spike sorting steps.

The recorded raw data should be filtered with a bandpass filter within the range of 300-6000 Hz to eliminate all the low- and high-frequency activities and highlight the spike shapes. This step is so critical and tricky as the whole spike shapes could be influenced by it. First, the filter should be non-causal to avoid any phase distortion. Second, the filter bandwidth should not be too narrow to prevent different missing features in spike shapes. After the filtering step, an amplitude threshold technique usually uses to detect spikes from the filtered data. Before extracting any features from the detected spikes, they must perfectly be aligned and sorted together. Then, they will be ready for the final clustering step by extracting the intended features from the aligned spikes.

It should be noted that choosing the feature is critical, as separating all the clusters depends on it. Thus, the extracted feature must have the ability to separate the spikes with the highest accuracy. Finally, as the last step, the spikes will be separated from each other based on similar features and form the clusters (3, 13). As an example, Wave_Clus (11) is an automatic and unsupervised spike sorting software that algorithm follows the same structure as mentioned before. After filtering the raw data and detecting the spikes by amplitude threshold technique, each spike’s wavelet coefficients are extracted and used as the input for the superparamagnetic clustering (SPC) (11). The open-access spike sorting algorithms with main features and specifications are listed as supplementary data (13). This study presents an interactive, offline, and unsupervised spike sorting software called ION-Decoding (Interactive Offline Neural Decoding) that can extract and align the spikes with a smart method also the clusters can be easily identified. The entire procedure is performed using the minimum number of adjustable parameters. Moreover, this study has a strong emphasis on PQRST assessment (14) in biomedical research. The software specifically designed for decoding single-unit activity from extracellular recording and have been successfully used on both simulated and human single-cell recording and the results were promising. It is presented in MATLAB and includes a userfriendly graphical interface. We will analyze each part of the presented algorithm (ION-Decoding) in detail.

The material section consists of information related to the benchmark dataset used for the algorithm assessment. Then, in the methods part, all the details associated with the algorithm’s different techniques and the statistical approaches are presented. After that, the results and outcomes of the algorithm are discussed. The discussion section is composed of a general review of the presented algorithm, and finally, all the examined information is summarized in the conclusion part.

## Material

The designed algorithm appraised by the simulated dataset produced from the database consists of 594 different average spikes collected from monkey basal ganglia and neocortex recordings to make the noise, multiunit, and single-unit activities(15). The sampling rate is 24 kHz, and there are 95 simulations with 10 minutes duration of extracellular recordings in the dataset. The number of single units varies between 2 to 20, and five different simulations were performed for each case. The data set is accessible from the webpage (http://bioweb.me/CPGJNM2012-dataset). The simulated data was generated based on the following steps (15):

As the initial step, the data was produced with a sample rate of 96000 Hz and then decimated by the factor of four for downsampling to 24000 Hz. The background noise simulation was achieved by modeling the overall contribution of neurons with a distance greater than 140 μm from the electrode tip via superimposing a large number of randomly selected average shape spikes from the database at random times. Then, the multiunit activity simulation was generated and added by a contribution of 20 different spike shapes chosen from the database to the background noise. The last step involved the simulation of single-unit activity caused by firing neurons close to the electrode tip. The firing times were generated with regards to a Poisson distribution with a mean firing rate randomly selected between 0.1 and 2 Hz, and the amplitude of each unit was randomly selected from a normal distribution within the range of 0.9 to 2 (15). This dataset has been used as a benchmark to assess different spike sorting algorithms’ performance in the literature(16–18).

## Methods

The block diagram of the designed algorithm is illustrated in **Fig. 2**.

**Fig. 2.**
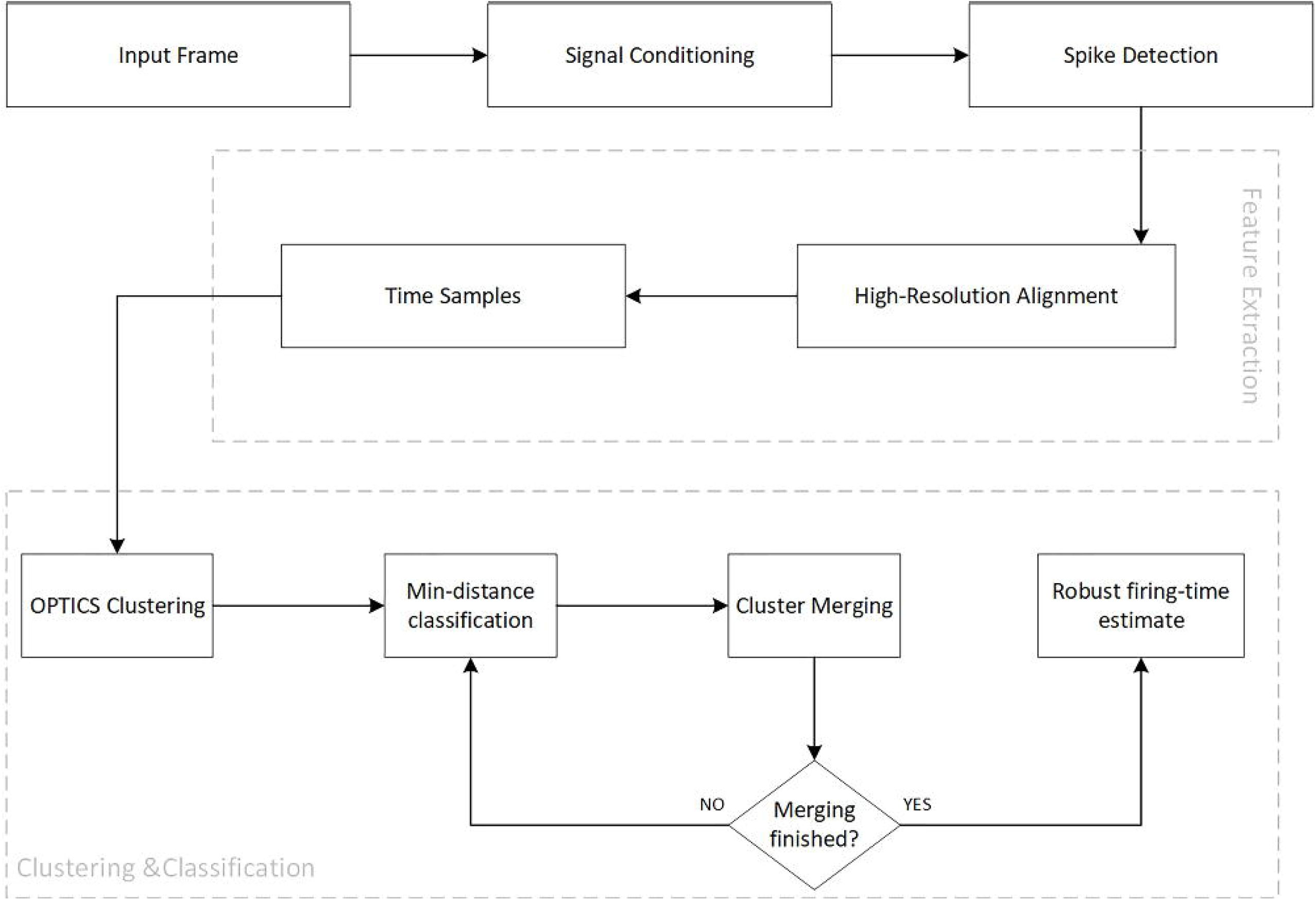
The main block scheme of the presented algorithm.

The details of this method are discussed in the following:

### Filtering

A four-pole digital bandpass Butterworth filter with the cut-off frequencies of 300 to 6000 Hz was used in the forward and reverse direction to prepare the recorded data and noise elimination. This filter has the advantage of reducing the Active Segments duration, decreasing temporal overlaps, and removing the low-frequency and non-discriminating elements from the signal (19).

### Spike Detection

After the filtering, the next step is the detection and extraction of spikes from the filtered data. This goal can be achieved by setting a threshold on the intended data. The purpose of the threshold setting is to exclude the spikes far from the recording electrode (passive segments) from those near the recording site (active segments). Therefore, an automatic amplitude threshold technique was used for spike detection(11):

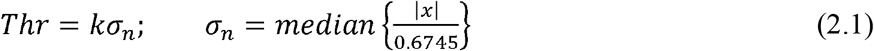

where *x* denotes the bandpass filtered data, *σ_n_* is an estimate of the standard deviation of the background noise, and *k* is the coefficient of the standard deviation that is usually set between 3 and 5. On the ION-Decoding algorithm, the parameter *k* was set to four, and active segments of 1ms intervals were extracted. This segment width makes it possible to mainly focus on the area near the signal’s peak to improve the quality of the algorithm when partial overlapping of the spikes occurs.

### Alignment and Feature Extraction

Feature extraction is usually performed in two parts. First, the signal has to align before entering the primary feature extraction step. To achieve this purpose, a high-resolution peak alignment method was used to align the detected active segments on the spikes’ highest peak. This method has the benefit of aligning and comparing waveforms and locating peaks (20). Briefly, the high-resolution alignment transfers the signal from discrete to the continuous domain (DFT interpolation) and then optimizes the phase shift by using Newton’s method. This method could align the active segments by sub-sampling intervals; thus, compensating for the problem raised by not sampling the AcS peaks during discretization.

### Clustering

The clustering method is implemented in this study called Ordering Points To Identify the Clustering Structure (OPTICS). OPTICS is a density-based clustering algorithm that is capable of computing an augmented cluster-ordering of the database objects. It can illustrate the density-based clustering structure of the database. The main advantage of this clustering algorithm rather than other methods (i.e., DBSCAN(21), K-mean(22), etc.) is that not only the method did not limit to one global parameter setting, but also the augmented cluster-ordering includes information which is equivalent to the density-based clustering’s corresponding to a broad range of parameter settings. It also supports non-convex clusters and clusters with different dispersions. As a result, this is a versatile basis for automatic and interactive cluster analysis (23). The clustering procedure is described as the following:

### Parameters and Definitions

The central concept of density-based clustering is that the surrounded neighboring objects included in a cluster should contain the minimum number of objects (23).

According to this description, the OPTICS algorithm works based on the two main input parameters (24):

- The scanning radius *ϵ* (the radius of the neighborhood of each object in a data set)
- *k*, the minimum number of objects considered as a cluster. The algorithm is designed based on some basic and core definitions introduced in Ref (23). However, the core distance and the reachability distance are the two main definitions needed to describe OPTICS.
- Core Distance (CD): A distance between the *ith* object and its k neighbor; i.e., each object has its individual core distance.
- Reachability Distance (RD): The RD of the *jth* object is the maximum distance between the *jth* object and the nearest object (d_ji_) and the core distance of the nearest object.

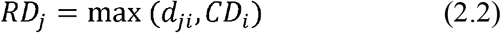 For using the OPTICS on the presented spike sorting algorithm, the *k* parameter is considered 50 empirically. However, the scanning radius *ϵ* was set infinity, so each object is considered a core object.

### The Clustering Operation

In the following, we will discuss the concepts and the operation of the OPTICS algorithm and figure out how the clustering is performed by this method.

The algorithm will choose an object randomly and assign an arbitrary and high value to its RD in the first step. This selected object will get the checked sign and will be shown on the origin of the reachability plot with the corresponding values, as the first object. The adjacent object, which sits next to the first one, should be detected on the second step. To achieve this goal, the distance between all the remaining objects will be calculated regarding the first object (the OPTICS algorithm will calculate all the distances based on the Euclidean distance). In the third step, the object with the minimum distance (i.e., the nearest object to the first random object) will be selected and shown on the reachability plot, and the y-axis value is set to its RD distance. That processed object will get the checked sign accordingly, and by this time, all the remained objects will be compared with regards to this current one. As the last step, the algorithm will back to the third step and going through the same process until all the objects get the checked sign. Finally, the clustering and calculated results will be shown on the reachability plot.

### Reachability Plot

OPTICS has the advantage of indicating the cluster-ordering of a data set graphically (23). This plot is representing the reachability distance versus the order of the objects (24).

It is possible to extract the details related to the detected clusters from this plot. **Fig. 3** shows the reachability plot of a dataset that comes from the output of the ION-Decoding algorithm. The signal name is “C_Easy1_noise01_short” and it consists of 3 neurons (downloadable from http://www.vis.caltech.edu/~rodri/Wave_clus/Simulator.zip) (25). As is shown in the plot, there are 2 points in which the RD’s value increased dramatically, compared to the previous points (the points with green stars and red arrows). These points make three valleys. The number of valleys indicates the number of neurons in the recorded signal. There are points with the lowest RD values representing the cluster agent (the blue interval on the zoomed figure; a.k.a., cluster representative). Any selected location amongst the lowest RD’s can be selected as the cluster agent. The clustering agent is the indicator of a cluster whereby all the remaining signals of the dataset will assign to them (the points with red stars and orange arrows).

**Fig. 3.**
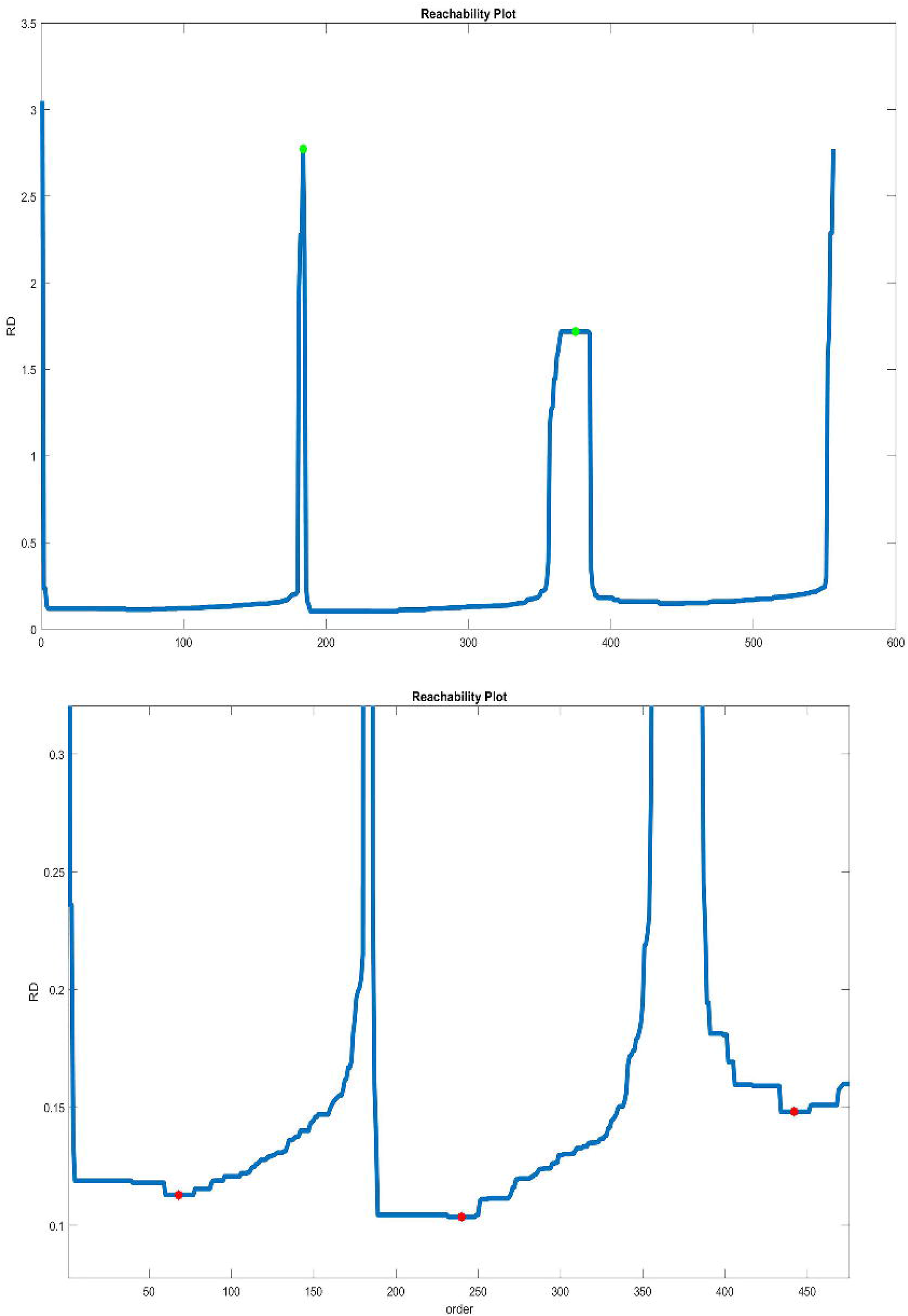
Examining the Reachability Plot from the OPTICS algorithm (up, the RD-plot; bottom, the zoomed version).

ION-Decoding algorithm can find these points (the cluster agents) automatically by identifying the local minimums of the RD-plot. Note that overestimating the number of clusters is preferred to its under sedation as it is possible to correct it by the “merging” step discussed later on.

### Classification

When the cluster agents (representative) are identified as the AcS’s corresponding with the valleys of the RD-plot, other segments are classified based on the following concepts:

The best two cluster representative is identified in terms of the Euclidean distance with the analyzed AcS and whether their correlation with the segment is higher than 0.8. Then, it is checked whether adding the new AcS creates firing time inconsistency (i.e., an IPI less than 2 ms) or not. The most similar cluster representative is selected if the above conditions are met.

Otherwise, the second cluster is selected. If both clusters do not meet the condition, no assignment is made.

### Merging

A more realistic RD-plot of the spike sorting is shown in **Fig. 4**. This kind of recordings usually includes a large number of neurons and spikes. As a result, the distance (RD) between the objects (the spikes) declines significantly, especially in dense datasets. This could cause problems in detecting pinnacle points (points with maximum RD between each valley) and cluster agents accordingly. Thus, a merging algorithm was designed to combine local points related to the same valleys.

**Fig. 4.**
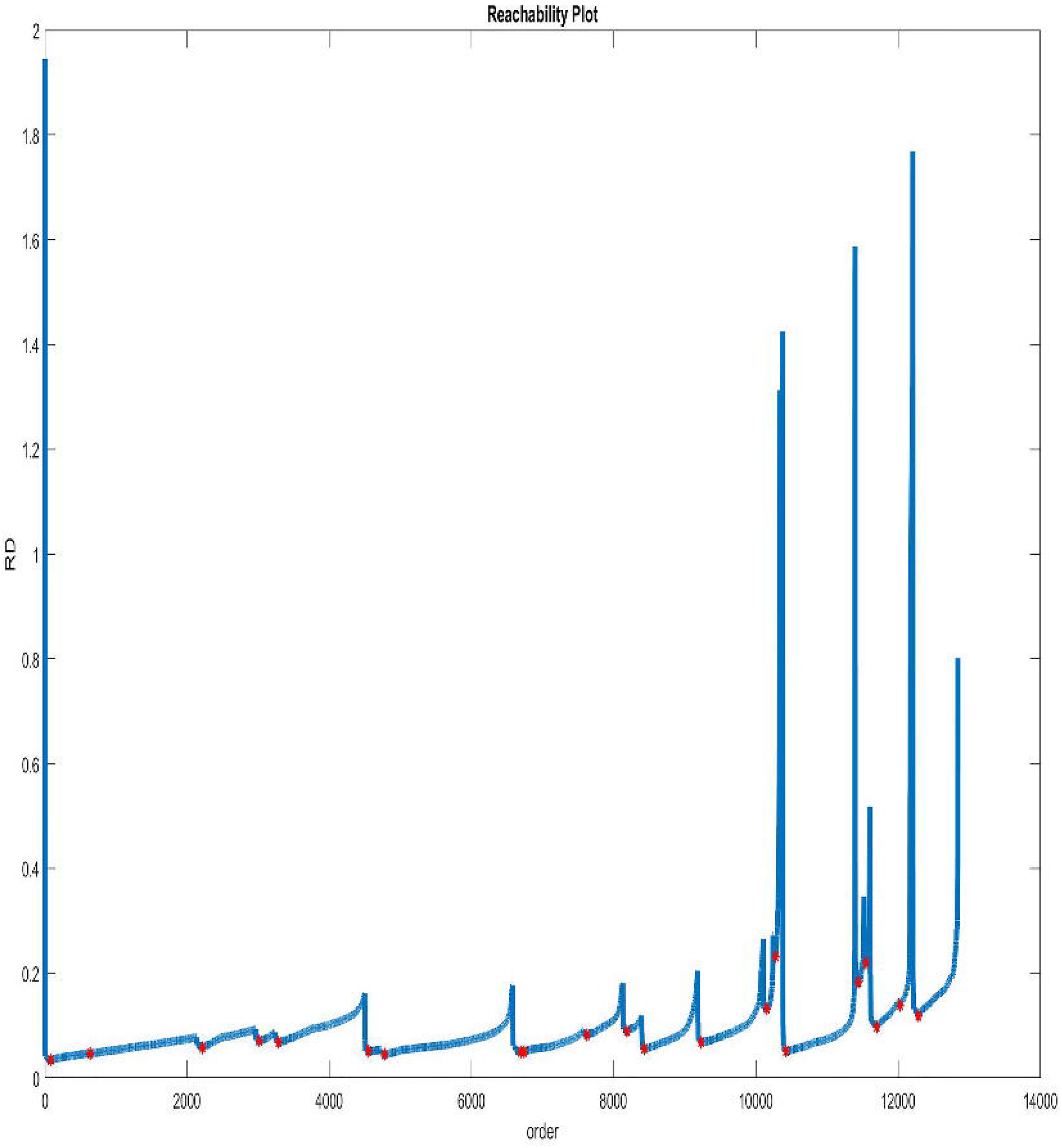
RD plot in a “simulation_95” dataset.

The merging algorithm is described as the following:

The correlation coefficient (CC) is calculated for two adjacent cluster agents. Moreover, the number of firing-time inconsistencies of the merged cluster (n_inc_)is calculated.

The two adjacent clusters are merged if the CC is higher than 0.8 (i.e., their cluster representatives are highly correlated) and n_inc_<=5. Such a tolerance in the firing-time inconsistency is accepted in the next step; we use the robust measure for neuron’s firing time estimation. Moreover, when decoding a large number of neurons, false-positive errors usually exists.

Also, the following conditions were considered to enhance the accuracy of the merging procedure:

### Robust firing-time estimate

To calculate the mean discharge rate (MDR) of each neuron, the trimmed mean (10%) of the Inter-Pulse-Interval (IPI) of the neurons are identified. The MDR is then estimated as the reciprocal of the trimmed mean value. We avoided using the simple averaging as it is not robust against unusually longer and shorter IPI’s when False Negative and Positive errors occur during classification. In the trimmed mean calculation, the IPIs are sorted, and then 5% of the higher and lower IPIs are removed, and then the average of the remained IPIs is calculated.

### Validation

The result of the automatic neural decoding (ION-decoding) was compared with that of the gold standard (a.k.a., the ground truth) in terms of the number of correctly identified neurons (clustering and merging) and also the accuracy of identified firings of each neuron (classification).

### Clustering accuracy

The clustering operation of the proposed algorithm was assessed by comparing the number of clusters in the gold standard (*n_g_*), the number of detected clusters by the algorithm (*n_m_*) and the number of common clusters between them (*n_c_*).

It is then possible to extract the number of missed and extra neurons based on the below equations:

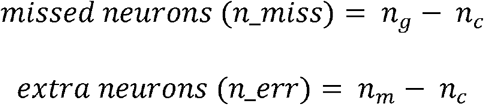

### Classification accuracy

The classification accuracy (a.k.a., agreement rate) of the algorithm was then evaluated for each common cluster (i.e., neuron) as below:

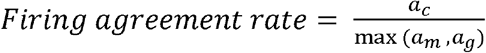

where *a_g_* is the number of spikes in a gold-standard neuron related, *a_m_* is the number of spikes in the corresponding neuron detected by the algorithm, and *a_c_* is the number of common spikes between the gold standard and algorithm output accordingly.

### Decomposability Index

The decomposability index (DI) parameter was calculated for each common cluster as the following:

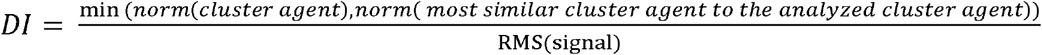

where RMS is the root-mean-square of the filtered signal. DI is very similar to the parameter signal-to-noise-ratio (SNR) used in communication that shows the complexity of decoding a specific neuron (26).

### Running time

The running time of the ION-Decoding algorithm for the entire 95 signals was calculated as a measure of computational complexity. The hardware specification of the computer is represented in table 1. The ION-Decoding program was written in MATLAB Release 2018b (MathWorks, Inc., Natick, Massachusetts, United States) and is available from the corresponding author to the interested readers.

**Table 1.**
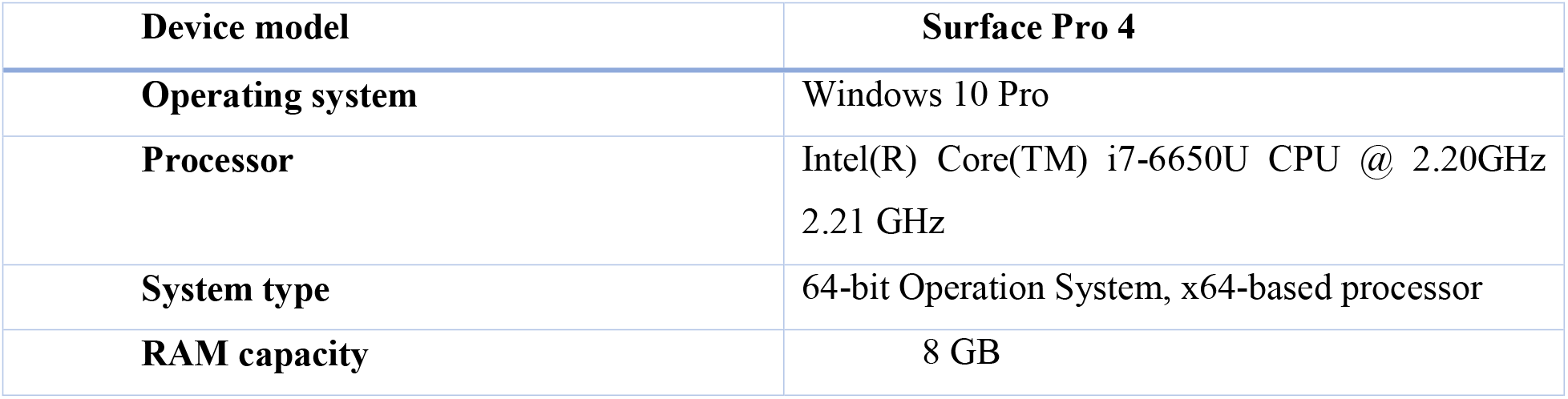
Hardware and software specifications

Results are reported as mean ± standard deviation. The normality of the estimated MDR and that of the corresponding gold standard was checked using the Kolmogorov-Smirnov (KS) test. When the normality was approved, the paired sample t-test was used to investigate whether the estimated MDR significantly differs from the gold standard in common units. Otherwise, the Wilcoxon signed-rank test was used. For examining the relationship between two continuous variables, the Pearson correlation coefficient was used when both variables were normally distributed. Otherwise, Spearman’s rank correlation coefficient was used. P-values less than 0.05 were considered significant. All statistical analyses and calculations were performed using the SPSS statistical package, version 18.0 (SPSS Inc., Chicago, IL, USA).

## Results and Discussion

The ION-Decoding algorithm was performed on one of the simulated datasets, simulation_1, as an example. It consists of 16 neurons with a duration of 10 minutes of extracellular recording. The SNR in this signal is about 40.6. The details of the signal and spikes shapes are represented in table 2 and Fig. 5, respectively.

**Fig. 5.**
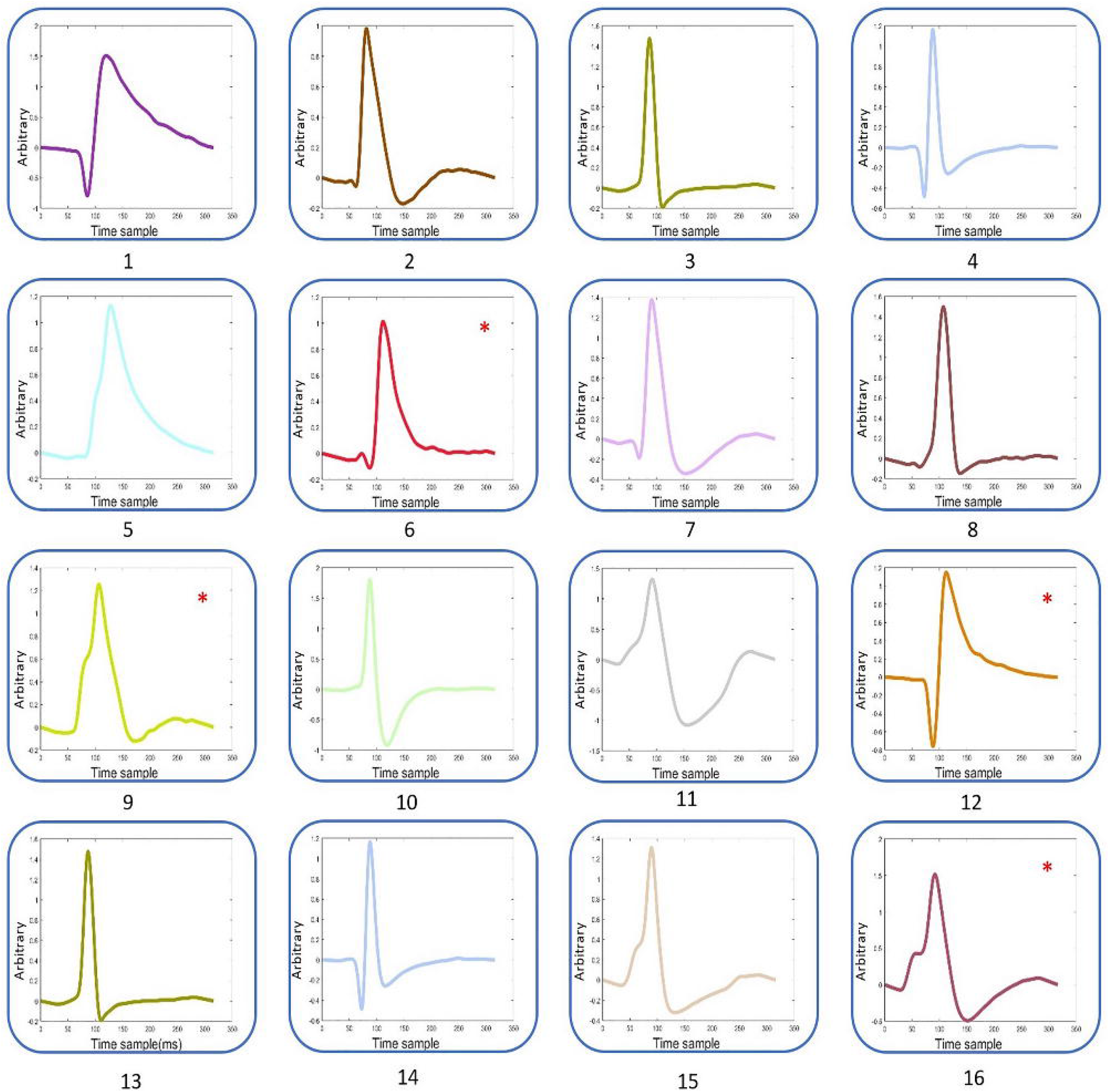
The Spikes shapes in the “simulation_1” dataset related to Table 2. Pictures with a “*” sign indicate those that could not be detected by ION-Decoding.

**Table 2.**
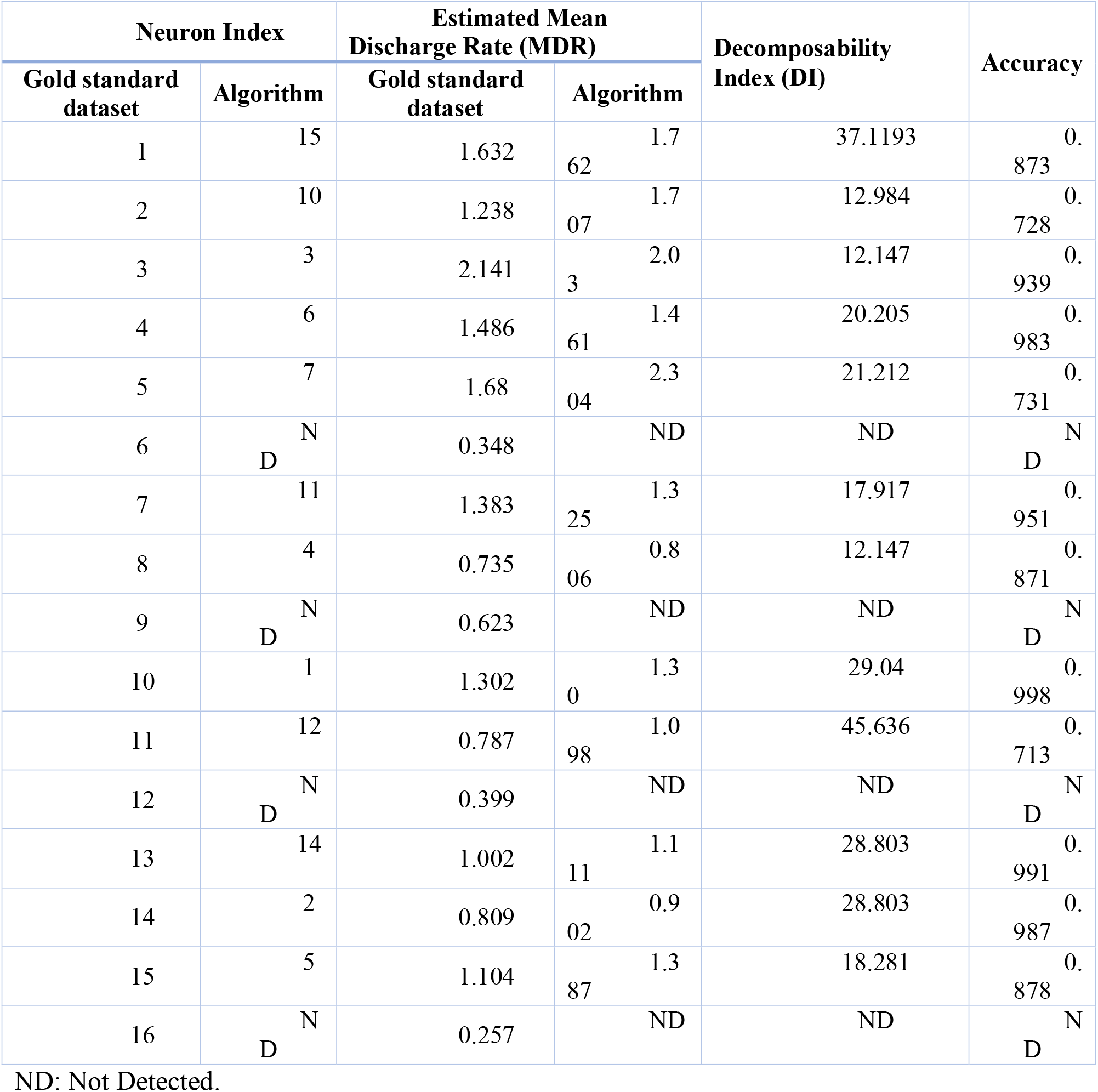
The result of the ION-Decoding on the simulation_1 signal.

Fig. 6 shows the GUI of the developed program. The output of the “simulation_1” was provided in this example. The ION-Decoding platform detected 15 neurons based on the RD plot, among which 12 neurons were correctly identified compared with 16 neurons in the gold standard. The histogram plot for the entire correctly detected neurons in Fig. 6 was shown in Fig 7.

**Fig. 6.**
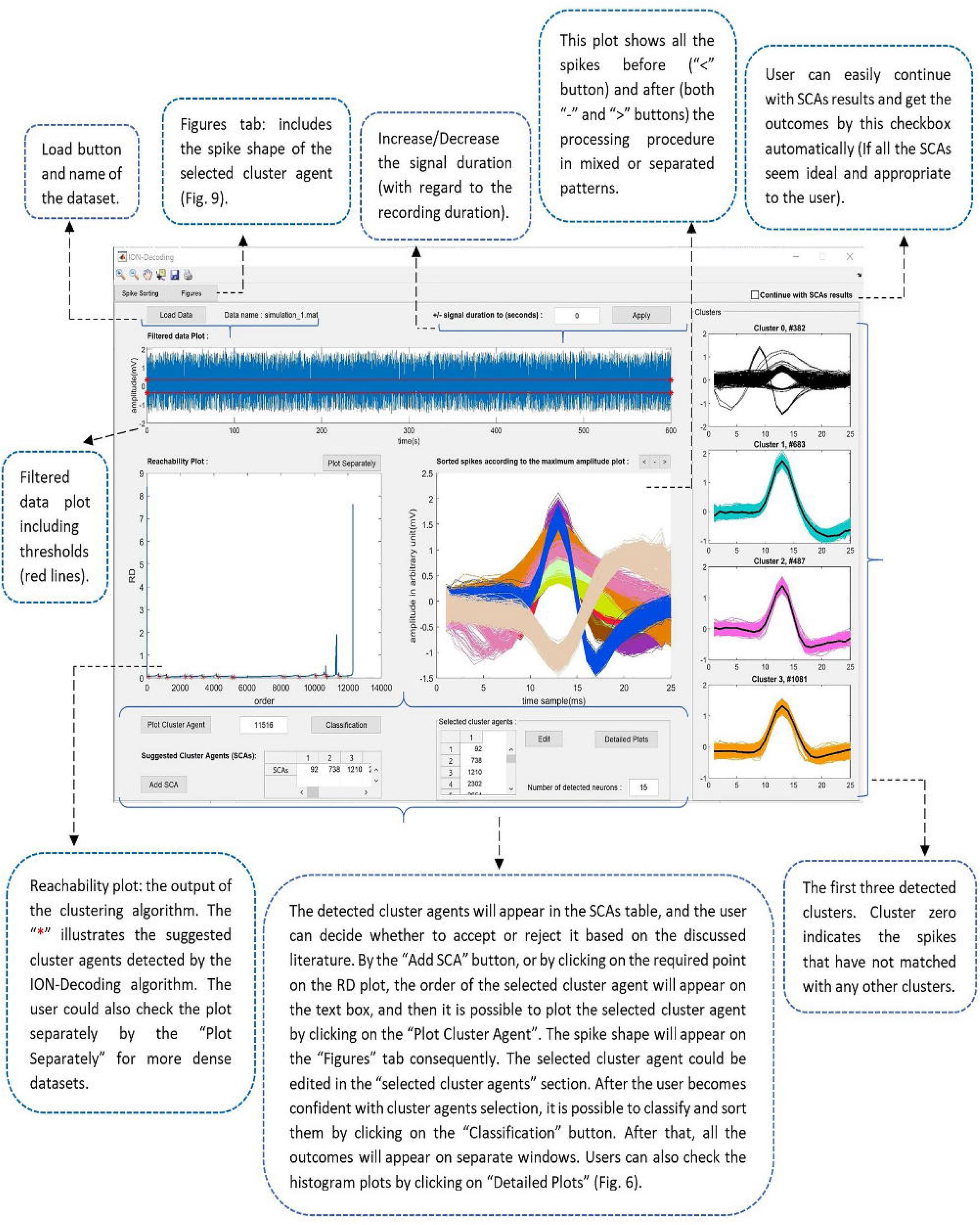
The GUI of the ION-Decoding algorithm for the dataset “simulation_1”.

**Fig. 7.**
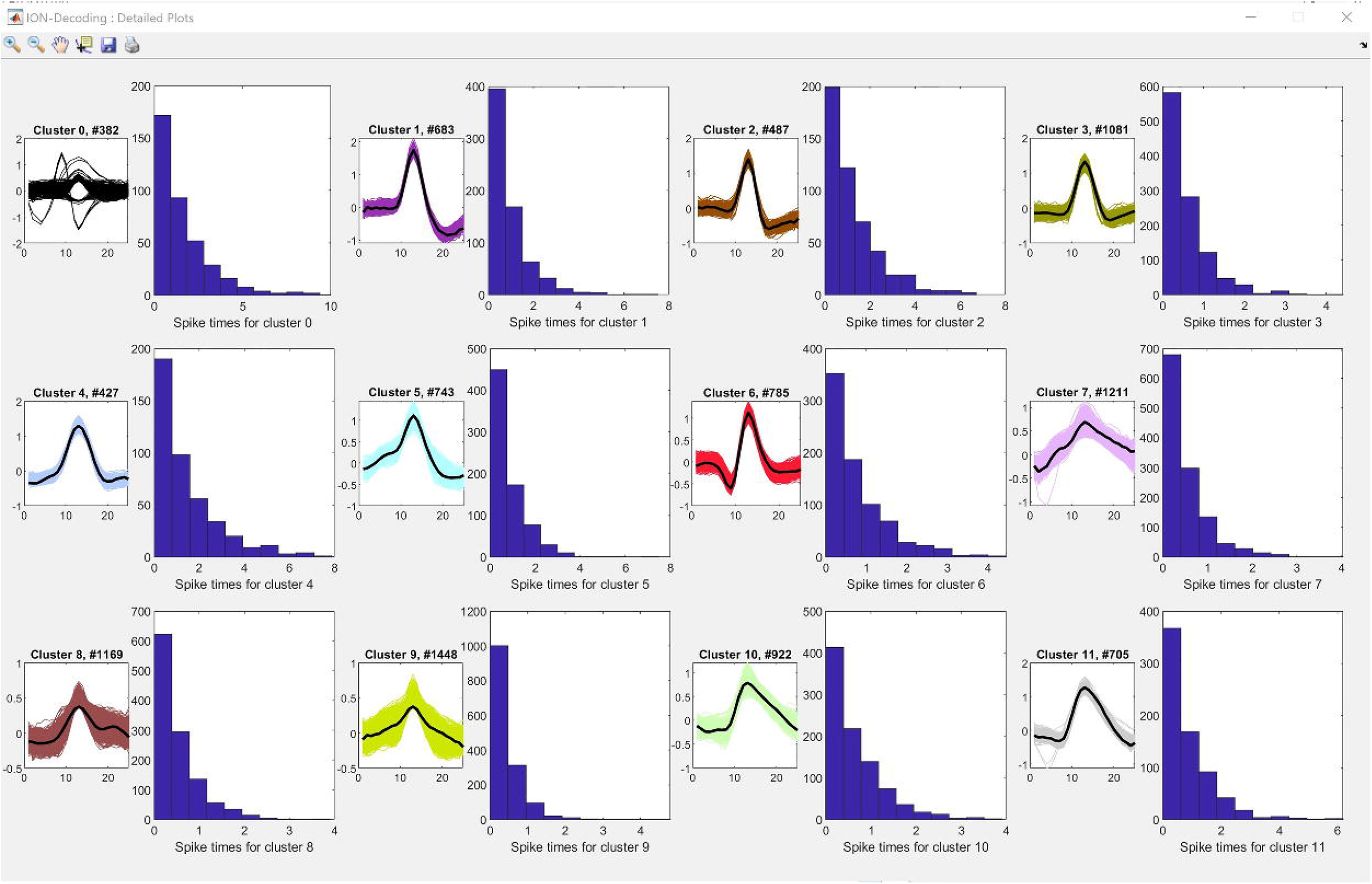
The histogram plots and spikes shape all the detected neurons by the ION-Decoding algorithm for the “simulation_1” dataset.

The average of the SNR of the entire 95 available signals in the dataset was (34.5±5.5). The performance of the clustering and classification procedures of the ION-Decoder was provided in Table 3.

**Table 3.**
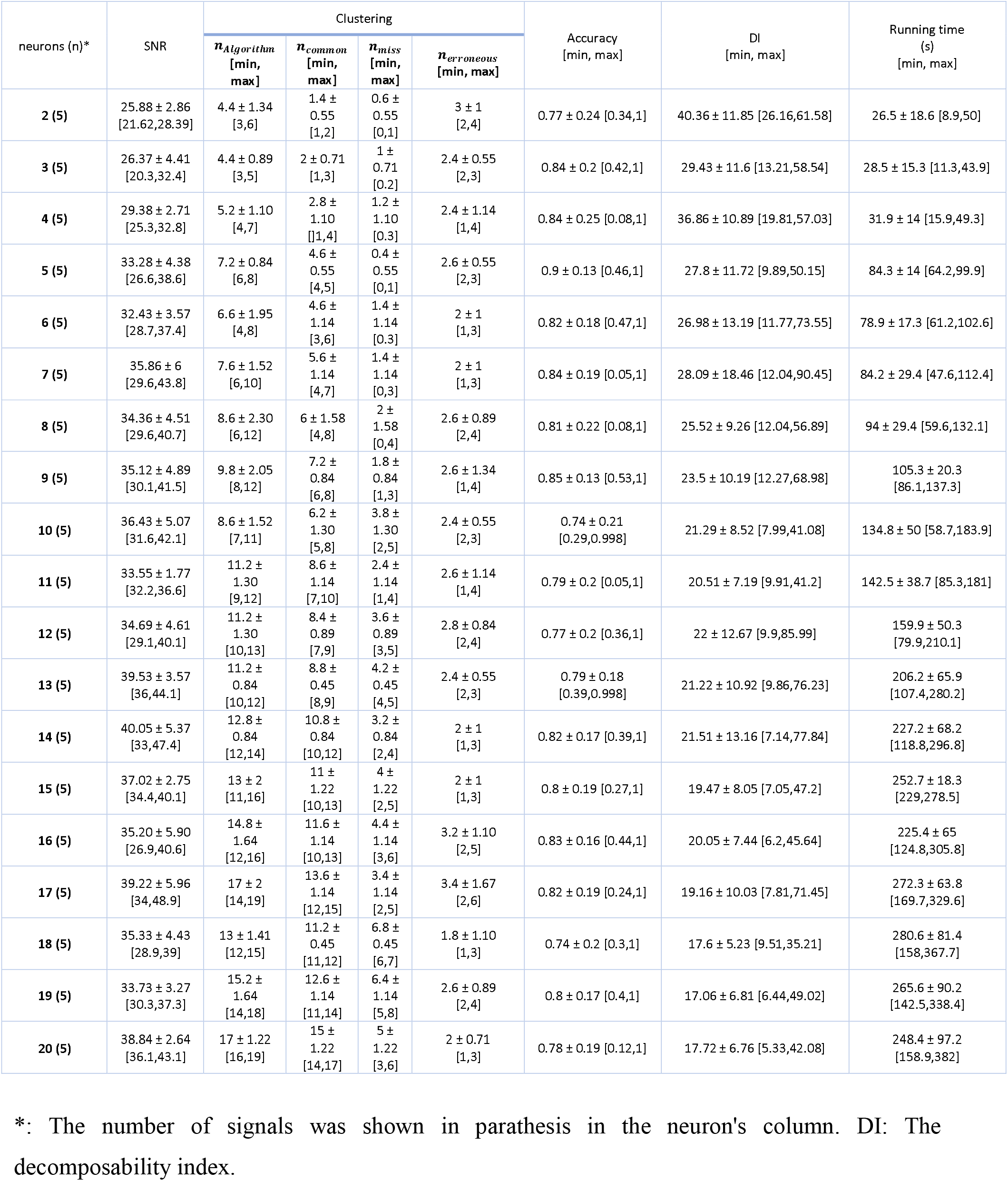
The performance of the ION-Decoding algorithm for the entire 95 signals.

| neurons (n)* | SNR | Clustering |  |  |  | Accuracy<br>[min, max] | DI<br>[min, max] | Running time<br>(s)<br>[min, max] |
| --- | --- | --- | --- | --- | --- | --- | --- | --- |
| | | $n_{Algorithm}$<br>[min, max] | $n_{common}$<br>[min, max] | $n_{miss}$<br>[min, max] | $n_{erroneous}$<br>[min, max] | | | |
| 2 (5) | $25.88 \pm 2.86$<br>[21.62, 28.39] | $4.4 \pm 1.34$<br>[3, 6] | $1.4 \pm 0.55$<br>[1, 2] | $0.6 \pm 0.55$<br>[0, 1] | $3 \pm 1$<br>[2, 4] | $0.77 \pm 0.24$ [0.34, 1] | $40.36 \pm 11.85$ [26.16, 61.58] | $26.5 \pm 18.6$ [8.9, 50] |
| 3 (5) | $26.37 \pm 4.41$<br>[20.3, 32.4] | $4.4 \pm 0.89$<br>[3, 5] | $2 \pm 0.71$<br>[1, 3] | $1 \pm 0.71$<br>[0, 2] | $2.4 \pm 0.55$<br>[2, 3] | $0.84 \pm 0.2$ [0.42, 1] | $29.43 \pm 11.6$ [13.21, 58.54] | $28.5 \pm 15.3$ [11.3, 43.9] |
| 4 (5) | $29.38 \pm 2.71$<br>[25.3, 32.8] | $5.2 \pm 1.10$<br>[4, 7] | $2.8 \pm 1.10$<br>[1, 4] | $1.2 \pm 1.10$<br>[0, 3] | $2.4 \pm 1.14$<br>[1, 4] | $0.84 \pm 0.25$ [0.08, 1] | $36.86 \pm 10.89$ [19.81, 57.03] | $31.9 \pm 14$ [15.9, 49.3] |
| 5 (5) | $33.28 \pm 4.38$<br>[26.6, 38.6] | $7.2 \pm 0.84$<br>[6, 8] | $4.6 \pm 0.55$<br>[4, 5] | $0.4 \pm 0.55$<br>[0, 1] | $2.6 \pm 0.55$<br>[2, 3] | $0.9 \pm 0.13$ [0.46, 1] | $27.8 \pm 11.72$ [9.89, 50.15] | $84.3 \pm 14$ [64.2, 99.9] |
| 6 (5) | $32.43 \pm 3.57$<br>[28.7, 37.4] | $6.6 \pm 1.95$<br>[4, 8] | $4.6 \pm 1.14$<br>[3, 6] | $1.4 \pm 1.14$<br>[0, 3] | $2 \pm 1$<br>[1, 3] | $0.82 \pm 0.18$ [0.47, 1] | $26.98 \pm 13.19$ [11.77, 73.55] | $78.9 \pm 17.3$ [61.2, 102.6] |
| 7 (5) | $35.86 \pm 6$<br>[29.6, 43.8] | $7.6 \pm 1.52$<br>[6, 10] | $5.6 \pm 1.14$<br>[4, 7] | $1.4 \pm 1.14$<br>[0, 3] | $2 \pm 1$<br>[1, 3] | $0.84 \pm 0.19$ [0.05, 1] | $28.09 \pm 18.46$ [12.04, 90.45] | $84.2 \pm 29.4$ [47.6, 112.4] |
| 8 (5) | 34.36 ± 4.51<br>[29.6,40.7] | 8.6 ± 2.30<br>[6,12] | 6 ± 1.58<br>[4,8] | 2 ± 1.58<br>[0,4] | 2.6 ± 0.89<br>[2,4] | 0.81 ± 0.22 [0.08,1] | 25.52 ± 9.26 [12.04,56.89] | 94 ± 29.4 [59.6,132.1] |
| 9 (5) | 35.12 ± 4.89<br>[30.1,41.5] | 9.8 ± 2.05<br>[8,12] | 7.2 ± 0.84<br>[6,8] | 1.8 ± 0.84<br>[1,3] | 2.6 ± 1.34<br>[1,4] | 0.85 ± 0.13 [0.53,1] | 23.5 ± 10.19 [12.27,68.98] | 105.3 ± 20.3<br>[86.1,137.3] |
| 10 (5) | 36.43 ± 5.07<br>[31.6,42.1] | 8.6 ± 1.52<br>[7,11] | 6.2 ± 1.30<br>[5,8] | 3.8 ± 1.30<br>[2,5] | 2.4 ± 0.55<br>[2,3] | 0.74 ± 0.21<br>[0.29,0.998] | 21.29 ± 8.52 [7.99,41.08] | 134.8 ± 50 [58.7,183.9] |
| 11 (5) | 33.55 ± 1.77<br>[32.2,36.6] | 11.2 ± 1.30<br>[9,12] | 8.6 ± 1.14<br>[7,10] | 2.4 ± 1.14<br>[1,4] | 2.6 ± 1.14<br>[1,4] | 0.79 ± 0.2 [0.05,1] | 20.51 ± 7.19 [9.91,41.2] | 142.5 ± 38.7 [85.3,181] |
| 12 (5) | 34.69 ± 4.61<br>[29.1,40.1] | 11.2 ± 1.30<br>[10,13] | 8.4 ± 0.89<br>[7,9] | 3.6 ± 0.89<br>[3,5] | 2.8 ± 0.84<br>[2,4] | 0.77 ± 0.2 [0.36,1] | 22 ± 12.67 [9.9,85.99] | 159.9 ± 50.3<br>[79.9,210.1] |
| 13 (5) | 39.53 ± 3.57<br>[36,44.1] | 11.2 ± 0.84<br>[10,12] | 8.8 ± 0.45<br>[8,9] | 4.2 ± 0.45<br>[4,5] | 2.4 ± 0.55<br>[2,3] | 0.79 ± 0.18<br>[0.39,0.998] | 21.22 ± 10.92 [9.86,76.23] | 206.2 ± 65.9<br>[107.4,280.2] |
| 14 (5) | 40.05 ± 5.37<br>[33,47.4] | 12.8 ± 0.84<br>[12,14] | 10.8 ± 0.84<br>[10,12] | 3.2 ± 0.84<br>[2,4] | 2 ± 1<br>[1,3] | 0.82 ± 0.17 [0.39,1] | 21.51 ± 13.16 [7.14,77.84] | 227.2 ± 68.2<br>[118.8,296.8] |
| 15 (5) | 37.02 ± 2.75<br>[34.4,40.1] | 13 ± 2<br>[11,16] | 11 ± 1.22<br>[10,13] | 4 ± 1.22<br>[2,5] | 2 ± 1<br>[1,3] | 0.8 ± 0.19 [0.27,1] | 19.47 ± 8.05 [7.05,47.2] | 252.7 ± 18.3<br>[229,278.5] |
| 16 (5) | 35.20 ± 5.90<br>[26.9,40.6] | 14.8 ± 1.64<br>[12,16] | 11.6 ± 1.14<br>[10,13] | 4.4 ± 1.14<br>[3,6] | 3.2 ± 1.10<br>[2,5] | 0.83 ± 0.16 [0.44,1] | 20.05 ± 7.44 [6.2,45.64] | 225.4 ± 65<br>[124.8,305.8] |
| 17 (5) | 39.22 ± 5.96<br>[34,48.9] | 17 ± 2<br>[14,19] | 13.6 ± 1.14<br>[12,15] | 3.4 ± 1.14<br>[2,5] | 3.4 ± 1.67<br>[2,6] | 0.82 ± 0.19 [0.24,1] | 19.16 ± 10.03 [7.81,71.45] | 272.3 ± 63.8<br>[169.7,329.6] |
| 18 (5) | 35.33 ± 4.43<br>[28.9,39] | 13 ± 1.41<br>[12,15] | 11.2 ± 0.45<br>[11,12] | 6.8 ± 0.45<br>[6,7] | 1.8 ± 1.10<br>[1,3] | 0.74 ± 0.2 [0.3,1] | 17.6 ± 5.23 [9.51,35.21] | 280.6 ± 81.4<br>[158,367.7] |
| 19 (5) | 33.73 ± 3.27<br>[30.3,37.3] | 15.2 ± 1.64<br>[14,18] | 12.6 ± 1.14<br>[11,14] | 6.4 ± 1.14<br>[5,8] | 2.6 ± 0.89<br>[2,4] | 0.8 ± 0.17 [0.4,1] | 17.06 ± 6.81 [6.44,49.02] | 265.6 ± 90.2<br>[142.5,338.4] |
| 20 (5) | 38.84 ± 2.64<br>[36.1,43.1] | 17 ± 1.22<br>[16,19] | 15 ± 1.22<br>[14,17] | 5 ± 1.22<br>[3,6] | 2 ± 0.71<br>[1,3] | 0.78 ± 0.19 [0.12,1] | 17.72 ± 6.76 [5.33,42.08] | 248.4 ± 97.2<br>[158.9,382] |

\*: The number of signals was shown in parathesis in the neuron's column. DI: The decomposability index.

### Statistical results

The number of neurons of the gold standard (n_g_) was significantly associated with the SNR of the signal (Spearman’s rho=0.510, p_value<0.001). The parameter ng was also significantly correlated with the number of missed neurons (n_miss_). Thus, the gold standard neurons were considered a confounding variable when the association between the parameters n_miss_ and SNR was examined. There was not any significant relation between *n_miss* with the SNR by controlling the *n_g_* parameter (correlation coefficient −0.169, p-value = 0.103). The parameter n_g_ was not significantly related to the number of erroneous neurons (n_err_) (Spearman’s rho=-0.054, p_value=0.604). Thus, a simple bivariate correlation was used to assess the association between n_err_ and SNR. The parameter n_err_ was not significantly dependent on the parameter SNR (Spearman’s rho=-0.087, p_value=0.400). Thus, the parameter SNR does not significantly affect the performance of the clustering algorithm.

The classification method’s accuracy was significantly associated with the parameter DI (Spearman’s rho=0.454, p_value<0.001). The scatter plot between the Accuracy and DI was shown in Fig. 8. Briefly, 77% of the neurons with the DI higher than 20 had the classification accuracy higher than 80%. Moreover, the intraclass correlation coefficient (ICC) between the estimated and gold standard MDR was 0.66 [CI 95%: 0.61-0.70].

**Fig. 8.**
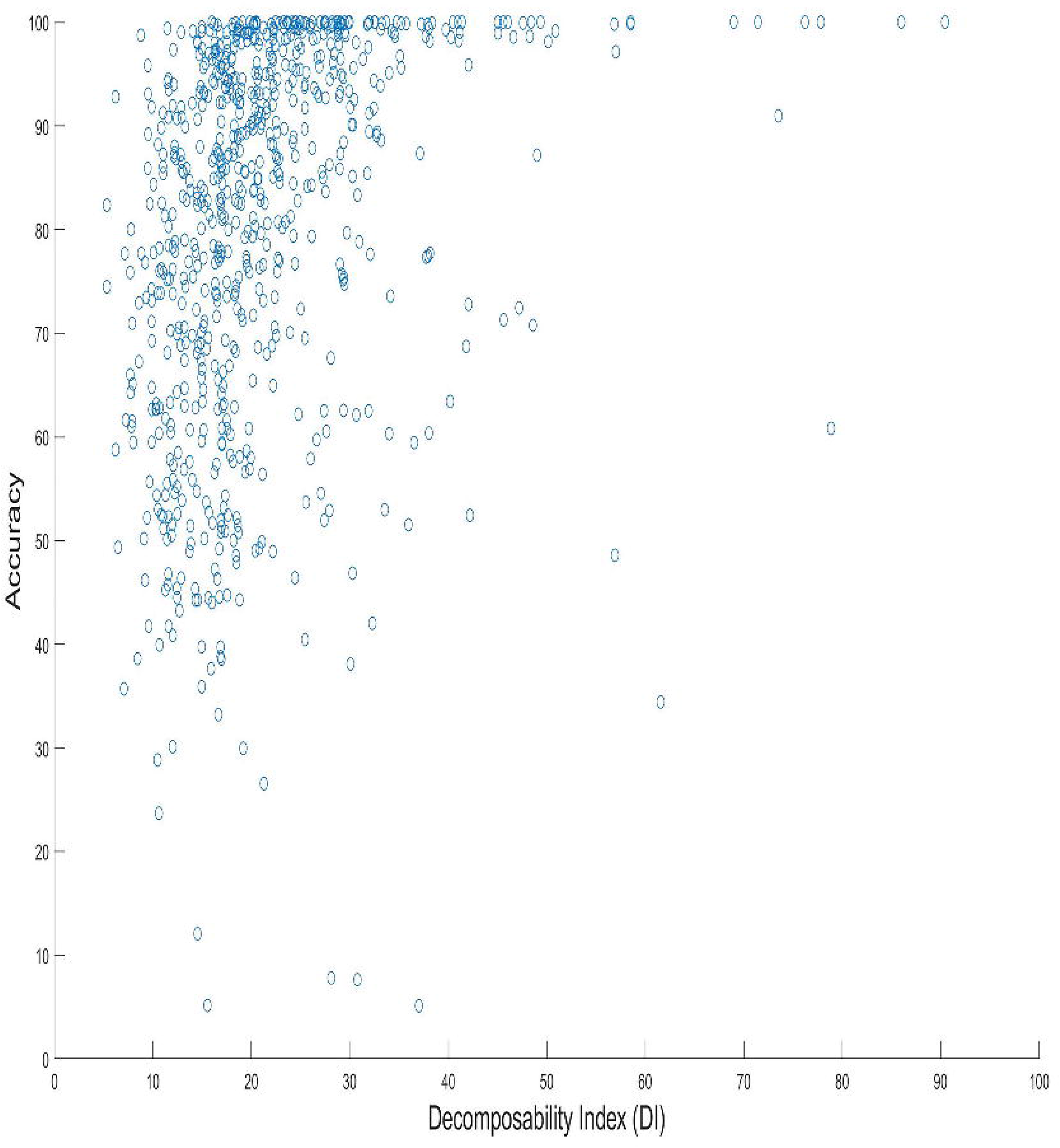
DI versus accuracy plot for all the common units in 95 database signals.

The performance of the proposed algorithm (ION-Decoder) was compared with the state-of-the-art (15). The number of hits (TP) and FP of ION-Decoder and Wave_Clus were compared in Table 4 on the benchmark dataset. The ION-Decoder significantly outperformed Wave_Clus in terms of the number of hits (p_value=0.017; Wilcoxon Signed Rank Test). However, The Wave_Clus algorithm significantly outperformed the ION-Decoding when FP error was considered (p_value=0.001; Wilcoxon Signed Rank Test).

**Table 4.** The comparison between the performance of the ION-Decoding and Wave_Clus

| Parameter | The number of hits |  |  |  |  |  | False Positive error |  |  |  |  |  |
| --- | --- | --- | --- | --- | --- | --- | --- | --- | --- | --- | --- | --- |
| Method | ION-Decoding |  |  | Wave_Clus |  |  | ION-Decoding |  |  | Wave_Clus |  |  |
| Neurons | Mean | Min | Max | Mean | Min | Max | Mean | Min | Max | Mean | Min | Max |
| 2 | 1.4 | 1 | 2 | 1.8 | 1 | 2 | 3.0 | 2 | 4 | 1.4 | 1 | 2 |
| 3 | 2.0 | 1 | 3 | 2.8 | 2 | 3 | 2.4 | 2 | 3 | 2.0 | 1 | 3 |
| 4 | 2.8 | 1 | 4 | 3.67 | 3 | 4 | 2.4 | 1 | 4 | 2.8 | 1 | 4 |
| 5 | 4.6 | 4 | 5 | 4.13 | 2 | 5 | 2.6 | 2 | 3 | 4.6 | 4 | 5 |
| 6 | 4.6 | 3 | 6 | 5.2 | 4 | 6 | 2.0 | 1 | 3 | 4.6 | 3 | 6 |
| 7 | 5.6 | 4 | 7 | 4.8 | 2 | 7 | 2.0 | 1 | 3 | 5.6 | 4 | 7 |
| 8 | 6.0 | 4 | 8 | 6.53 | 6 | 7 | 2.6 | 2 | 4 | 6.0 | 4 | 8 |
| 9 | 7.2 | 6 | 8 | 7 | 4 | 9 | 2.6 | 1 | 4 | 7.2 | 6 | 8 |
| 10 | 6.2 | 5 | 8 | 6.4 | 3 | 9 | 2.4 | 2 | 3 | 6.2 | 5 | 8 |
| 11 | 8.6 | 7 | 10 | 7.8 | 6 | 10 | 2.6 | 1 | 4 | 8.6 | 7 | 10 |
| 12 | 8.4 | 7 | 9 | 7.6 | 6 | 11 | 2.8 | 2 | 4 | 8.4 | 7 | 9 |
| 13 | 8.8 | 8 | 9 | 7.93 | 3 | 11 | 2.4 | 2 | 3 | 8.8 | 8 | 9 |
| 14 | 10.8 | 10 | 12 | 6.93 | 4 | 11 | 2.0 | 1 | 3 | 10.8 | 10 | 12 |
| 15 | 11.0 | 10 | 13 | 7.93 | 5 | 11 | 2.0 | 1 | 3 | 11.0 | 10 | 13 |
| 16 | 11.6 | 10 | 13 | 10.33 | 8 | 12 | 3.2 | 2 | 5 | 11.6 | 10 | 13 |
| 17 | 13.6 | 12 | 15 | 6.53 | 4 | 9 | 3.4 | 2 | 6 | 13.6 | 12 | 15 |
| 18 | 11.2 | 11 | 12 | 8.6 | 4 | 13 | 1.8 | 1 | 3 | 11.2 | 11 | 12 |
| 19 | 12.6 | 11 | 14 | 7.66 | 5 | 11 | 2.6 | 2 | 4 | 12.6 | 11 | 14 |
| 20 | 15.0 | 14 | 17 | 8.33 | 5 | 12 | 2.0 | 1 | 3 | 15.0 | 14 | 17 |

The ION-Decoding is thus a promising single-channel spike sorting algorithm. However, our future focuses on the improvement of the cluster representative identification and FP error reduction.

